# PI3K inhibition reverses migratory direction of single cells but not cell groups in electric field

**DOI:** 10.1101/2020.08.05.238170

**Authors:** Y Sun, H Yue, C Copos, K Zhu, Y Zhang, Y Sun, X Gao, B Reid, F Lin, M Zhao, A Mogilner

## Abstract

Motile cells migrate directionally in the electric field in a process known as galvanotaxis. Galvanotaxis is important in wound healing, development, cell division, and nerve growth. Different cell types migrate in opposite directions in electric fields, to either cathode, or anode, and the same cell can switch the directionality depending on chemical conditions. We previously reported that individual fish keratocyte cells sense electric fields and migrate to the cathode, while inhibition of PI3K reverses single cells to the anode. Many physiological processes rely on collective, not individual, cell migration, so here we report on directional migration of cohesive cell groups in electric fields. Uninhibited cell groups of any size move to the cathode, with speed decreasing and directionality increasing with the group size. Surprisingly, large groups of PI3K-inhibited cells move to the cathode, in the direction opposite to that of individual cells, which move to the anode, while such small groups are not persistently directional. In the large groups, cells’ velocities are distributed unevenly: the fastest cells are at the front of the uninhibited groups, but at the middle and rear of the PI3K-inhibited groups. Our results are most consistent with the hypothesis, supported by the computational model, that cells inside and at the edge of the groups interpret directional signals differently. Namely, cells in the group interior are directed to the cathode independently of their chemical state. Meanwhile, edge cells behave like the individual cells: they are directed to the cathode/anode in uninhibited/PI3K-inhibited groups, respectively. As a result, all cells drive uninhibited groups to the cathode, but a mechanical tug-of-war between the inner and edge cells directs large PI3K-inhibited groups with cell majority in the interior to the cathode, while rendering small groups non-directional.

**Significance statement:** Motile cells migrate directionally in electric fields. This behavior – galvanotaxis – is important in many physiological phenomena. Individual fish keratocytes migrate to the cathode, while inhibition of PI3K reverses single cells to the anode. Uninhibited cell groups move to the cathode. Surprisingly, large groups of PI3K-inhibited cells also move to the cathode, in the direction opposite to that of individual cells. The fastest cells are at the front of the uninhibited groups, but at the middle and rear of the PI3K-inhibited groups. We posit that inner and edge cells interpret directional signals differently, and that a tug-of-war between the edge and inner cells directs the cell groups. These results shed light on general principles of collective cell migration.

## INTRODUCTION

Cells migrate collectively, as cohesive groups, in development, wound healing, and tumor invasion [1, 2], yet our knowledge of cell migration comes largely from single-cell studies. An open fundamental biological problem is to understand the coordinated movement and directionality of cohesive cell groups. Experimental research [1, 2] on and modeling [3-6] of groups of cells migrating in chemical gradients brought much insight into mechanics and directional cell behavior. A few conceptual models emerged from this research, including, but not restricted to: A) There exist leader cells [7, 8], which are usually located at the group’s leading edge and are polarized and actively migrating in the direction of an external cue. The remaining cells follow the leader cells passively [5]. B) Inner cells in the group polarize in the direction of the external cue and migrate actively, while edge cells do not respond to the directional signal and are dragged and pushed along by the inner cells [6, 9]. C) The group is integrated mechanically [10], so that all cells are tightly interlinked into a supracellular tissue by cytoskeletal structures spanning the whole group [11].

Most of research on collective cell migration was done on groups moving in chemical gradients, yet there are other physiological directional cues that cells encounter. One of them is direct-current electric field (EF) guiding adhesive cell groups in development [12], wound healing [13], and regeneration [14]. Some types of cells (i.e. keratinocytes) individually migrate to the cathode (minus end) in EFs, others (i.e. fibroblasts) – to anode (plus end) [15]. Respective galvanotactic signals may be as potent as, or even more important than, chemotactic signals [16]. Electrically sensitive cells is the rule, not the exception [16]. Most of research on galvanotaxis (also called electrotaxis) was done on single cells, but a few recent studies started to investigate collective migration in EF of epithelial cell sheets [17, 18], MDCK cell groups [9, 19], corneal epithelial cells [20], and HaCaT cellular monolayers [21].

One cell type, fish epidermal keratocyte, has been instrumental in studying mechanisms of galvanotaxis due to its fast and steady locomotion, simple shape, and well-understood motile mechanics [22]. It has long since been known that keratocytes sense EF and move to the cathode [23]. Physically, keratocytes likely sense EF by harnessing electrophoresis of charged mobile transmembrane proteins, which aggregate to one of the cell’s sides in EF [24] and serve either as receptors activating intracellular signaling relays, or as scaffolds for such receptors. Inhibition of PI3-Kinase (PI3K, for brevity) redirects these cells to the anode [24, 25]. Collective keratocyte migration has received much less attention than single cell migration; however, both spontaneous migration of groups of zebrafish [26] and gold fish [3] keratocytes, and EF-induced collective movements of zebrafish [27] and gold fish [23] keratocytes have been characterized.

Many questions about cell groups migrating in EF remain unanswered: Do the cells sense EF individually and independently, or collectively? Are there leader and follower cells? Considering that some perturbations can reverse individual cell directionality, what would such perturbations do to the collective directionality? In this paper, we investigate movements of cohesive keratocyte groups in EF. We find that, expectedly, unperturbed keratocyte groups migrate to the cathode, same as individual cells. Surprisingly, large cohesive groups of PI3K-inhibited cells move to the cathode, opposite of the anode-directed single PI3K-inhibited keratocytes. Speed of cells inside the large unperturbed and PI3K-inhibited groups is maximal at the groups’ fronts and rears, respectively. These behaviors are consistent with a model according to which cells at the group edge sense EF in a similar way to the individual cells and tend to go to the cathode or anode depending on their chemical state (unperturbed or PI3K-inhibited). Meanwhile, cells in the group’s interior are not passive followers, but rather tend to polarize and move to the cathode, independently of their chemical state. In unperturbed groups, both inner and

edge cells are driven to the cathode, and the whole group moves to the cathode. A tug-of-war between the inner and edge cells in PI3K-inhibited groups determines the group’s directionality: in large groups, the inner cell majority drive the group to the cathode; small groups are not persistently directional.

## METHODS

### Primary culture of keratocyte groups and single cells

Adult zebrafish (strain AB), were obtained from the UC Davis Zebrafish facility. All experiments were conducted in accordance with the UC Davis Institutional Animal Use and Care Committee protocol 16478. Scales were removed from zebrafish flanks and allowed to adhere to the bottom of EF chambers [25, 28]. The scales were covered by a glass 22-mm coverslip with a stainless steel nut on the top, and cultured at room temperature in Leibovitz’s L-15 media (Gibco BRL), supplemented with 14.2 mM HEPES pH 7.4, 10% Fetal Bovine Serum (Invitrogen), and 1% antibiotic-antimycotic (Gibco BRL). Scales were removed gently once an epithelial sheet forms, which usually takes 24-48 hours. Cell groups of various sizes and shapes separated from the epithelial sheet spontaneously and migrated away; these groups were used in the experiments with EF. To isolate single cells, groups of keratocytes that migrated off the scale were digested by a brief treatment with 0.25% Trypsin/0.02 EDTA solution (Invitrogen) in phosphate buffered saline (PBS), and then kept on ice until use [28, 29].

### Pharmacological perturbation experiments

Drugs (all purchased from Sigma) were added in the culture medium in the following concentration: DMSO as drug control (0.1%), LY294002 (50 or 100 μM), Blebbistatin (50 μM), EGTA (5 mM). Subsequent experiments were implemented in the presence of each of these drugs within 15 minutes of incubation.

### EF application and time-lapse recording

The electric fields were applied as previously described [30, 31] in custom-made electrotaxis chambers to minimize heating during experiment. To eliminate toxic products from the electrodes that might be harmful to cells, agar salt bridges made with 1% agar gel in Steinberg’s salt solution were used to connect silver/silver chloride electrodes in beakers of Steinberg’s salt solution to pools of excess medium at either side of the chamber (Fig. S1A). EF strength was chosen based on physiological range and previous studies [25]. In most experiments, an EF of 1 V/cm was used unless otherwise indicated. The actual voltage is measured by a voltmeter before and after each experiment. Phase contrast images were captured by a Zeiss Observer Z1 (Carl Zeiss) equipped with a QuantEM:512SC EMCCD camera (Photometrics). Time-lapse experiments were performed using MetaMorph NX software controlling a motorized scanning stage (Carl Zeiss). Typically, in each experiment, overlapped fields covering a whole cell group were captured sequentially. Images were taken at 30 or 60 second interval at room temperature for up to 60 minutes unless stated otherwise.

### Image processing and data analysis/presentation

Time-lapse images were imported into ImageJ and stitched by using Grid/Collection Stitching plugin (Fig. S1B). To quantify single-cell and collective group motility, we extracted the trajectory of each object using an automatic/manual tracking tool [25, 32, 33] (Fig. S1C, zonal analysis). Directedness was defined as cos(*θ*), where *θ* is the angle the EF vector and the vector connecting the centroids of the cell/group in the initial and final positions [34, 35]. To quantify cell/group migration, each cell was numbered (Fig. S1E). x and y coordinates of single cells and of the groups’ centroid were measured in each image (the origin (0,0) was the initial coordinate) in the image stack, with the x-axis parallel to the EF, as described previously [25]. The (x, y) trajectory data were imported into ImageJ chemotaxis tool plugin and rescaled to physical dimensions based on pixel sizes.

### Aligning contours and mapping edge velocity

Serials polygonal outlines of a cell group were extracted from time-lapse images and sampled at 200 evenly spaced points. These contours were then mutually and sequentially aligned to simulate collective cell motion over time [36]. For the first frame of a time-lapse sequence, the contour was adjusted manually to make sure that the first point is located right in the middle facing the cathode. The cell sheet boundary positions were translated to polar coordinates. The edge velocity at each point was calculated by dividing the displacement vector normal to the cell group edge by the time interval [37]. A custom scalar map function written in Matlab was used to generate continuous space-time plots of protrusion and retraction (Fig. S1D).

### Morphological characterization and quantification

Stitched images were converted into binary images using custom written Matlab codes. Briefly, we used Matlab edge detection and a basic morphology function to outline cell groups in the phase contrast image. In most cases, we had to use the Lasso tool in Photoshop (Adobe) to manually draw the group shape (Fig. S1C, Global analysis). Polygonal outlines extracted from the binary images were plotted in Celltool, an open source software [36]. Geometric features of each cell group including centroid and area were measured directly from the polygons by using standard formulas [38]. Cell numbers (both edge and inner cells) were either manually counted (for the groups with less than 300 cells) using ImageJ particle analysis tool (Fig. S1E) or calculated based on the area fractions (for the groups with more than 300 cells).

### PIV analysis

Collective migration in the time-lapse images was quantified by PIV analysis using custom MATLAB code based on MatPIV1.6.1 (Fig. S1C, PIV analysis), as previously described [20]. Multiple iterations of interrogation window sizes were used (two iterations of 64 × 64 pixel windows followed by two iterations of 32 × 32 pixel windows). At each interrogation step, a 50% overlap was used; outliers were detected using a signal-to-noise filter. The resulting PIV vectors capture all motion within the cell group and displacements on a time scale of 1 minute. Kymographs were used to quantify and visualize spatiotemporal dynamics of the velocity component parallel to EF and of directedness. For each data matrix from the PIV analysis, we computed the average value for each column parallel to EF and then derived a one-dimensional segment for each time point. We also computed the time average of the velocity component parallel to EF and of directedness of each vector in each cell group during 10-min time period. Spatial variations of the directedness and velocity were visualized by color-coded surface plots.

### Statistics and reproducibility

All experiments were repeated and produced similar results. In most cases a representative experiment is shown, unless stated otherwise. Data are presented as means ± standard error. To compare group differences (EF vs no EF or drug treatment vs no treatment), either chi-squared test, or paired/unpaired, two-tailed Student’s t-test were used. A p value less than 0.05 was considered as significant.

### Modeling

We simulate the movement of the cells and cell groups using Cellular Potts Model (CPM) [39]. In our CPM implementation, we have a two-dimensional square lattice with each site (pixel) occupied by an integer ‘spin’, *s*. Each biological cell is represented by the sets of pixels with the same (non-zero) spin, while the substratum not covered by cell(s) is represented by the pixels with spin 0. In our model, the cells are initiated as 5×5 (25 pixels total) squares. The shape and position of the cells evolve by changing the spins of the pixels via a Monte Carlo algorithm based on the effective mechanical energy of the system. More specifically, at each computational step, the spin of one pixel can be changed to be equal to that of a randomly chosen neighboring (in the Moore neighborhood) pixel with a different spin based on the following probability:

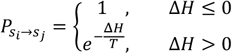

Here, *s*_*i*_ is the original spin of pixel *i* and *s*_*j*_ is the target spin copied from the chosen neighbor *j*. Δ*H* is the change of the effective energy if this change of spin is accepted, and *T* is the effective temperature parameter reflecting the cell edge fluctuation amplitude.

The effective energy is defined as:

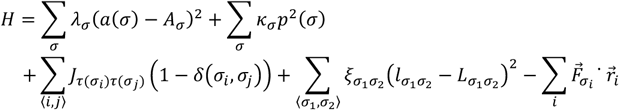

In this equation, the first two terms sum over all the cells with the first term representing the energy related to the intracellular pressure caused by cell area conservation, and the second term representing the energy related to the effective cell surface tension. More specifically, *σ*is the cell index, and the summation goes over all cells. Variables *a*(*σ*) and *p*(*σ*) are the area and perimeter of cell *σ*, and *A*_*σ*_ is the ‘target area’ of this cell. Parameters *λ*_*σ*_ and *k*_*σ*_ are responsible for the strength of the effects of conserving the area and perimeter of the cell. The third term is the effective adhesion energy between different cells; the summation goes over all pairs of neighboring pixels. Parameter 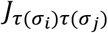 reflects the adhesion strength and depends on whether the interacting pair of pixels belong to different cells, or one to cells, and the other to the empty substratum. This term is non-zero only when *σ*_*i*_ and *σ*_*j*_ are two different cells (due to the term with delta-function), or a cell and the substratum. The fourth term is an effective elastic coupling between neighboring cells *σ*_1_ and *σ*_2_. Variable 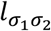 is the distance between the centroids of these two cells, *ξ* is the spring constant, and parameter 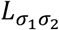 is the target distance. This elastic energy only exists between neighboring cells with cell-cell distance smaller than a threshold. The last term is the effective potential energy designed to move the cells in the direction of their polarization by applying forces to them. The driving forces 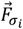 for different cells can have different strengths and directions depending on cells’ chemical states and positions within the group. 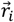 is the position of pixel *i*. We explored two models in which the parameters *λ, κ, J* are held constant but the driving force in the last term is varied, as explained below. We use CompuCell3D [40] for the numerical simulations. Parameters values in the models are non-dimensional and are listed in the Supplemental Table.

The key part of the models is the magnitude and direction of the driving forces 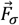. To match the experiment, we include noise to the force’s direction (in addition to the inherent thermal noise of the cells’ edges in CPM). The equation for the evolution of one cell’s direction of the driving force) is:

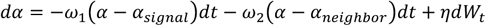

In this equation, *α* is the cell’s polarization direction (angle in radians with respect to the direction of the cathode), which is the same as the direction of vector 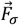 The first term means that the cell tends to align its polarizing direction to the EF signal direction that it senses. For example, for the uninhibited case, the cells tend to move towards the cathode so that *α*_*signal*_ = 0, while for the PI3K-inhibited case, the inner cells still tend to move to the cathode with *α*_*signal*_ = 0, while the edge cells tend to move towards the anode so that *α*_*signal*_ = *π*. The second term means that the cell tends to align with its neighbors. This term does not exist for edge cells in the PI3K-inhibited case, as we assume that for this case, some communications between the edge cells and inner cells are lost. To calculate *α*_*neighbor*_, we separate the neighbor cells’ influence into two directions: x and y direction as follows: *Eff*_*x*_ = ∑_*i,neighbor*_ cos(*α*_*i*_) and *Eff*_*y*_ = ∑_*i,neighbor*_ sin(*α*_*i*_). Then, total effect 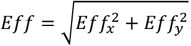. So, 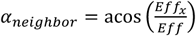 when *Eff*_*y*_ > 0 and 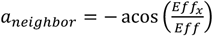 when *Eff*_*y*_ < 0 Parameters *ω*_1_ and *ω*_2_ are responsible for the alignment rates. The third term is the stochastic noise with parameter *η* responsible for the noise amplitude, and *dW*_*t*_ denoting the Wiener process (uncorrelated white noise). During each step in the simulation, Δ*W*_*t*_ *≡ W*_*t+1*_ *W*_*t*_∼*N*(0,1), with *N*(0,1) denoting the standard normal distribution.

After the polarization direction *α* is determined from the above equations, the driving forces for the cell can be determined as follows. For all cells except the second layer of cells in the group (inner cells that have edge cells as nearest neighbors), the driving forces just follow the polarizing direction: *F*_*x*_ = *Fcosα* and *F* _*y*_= *Fsinα*. (*F* can be *F*_*inner*_ or *F*_*edge*_ depending on the cell’s position.) In order to soften drastic possible collisions between the edge cells and inner cells closest to them and driven in opposite direction, we add the following to the model: For the second layer of cells, in both uninhibited and PI3K-inhibited cases, the cells are influenced by the edge cells, but only when the edge cells move inward, toward the second layer cells. More specifically, for any second layer cell, *F*_*x*_ = *F*_*inner*_*cosα + ∑*_*i,edge*_*inward*_ *F*_*i*_*cosβcosγ, F*_*y*_ = *F*_*inner*_*sinα + ∑*_*i,edge*_*inward*_ *F*_*i*_ *cosβsinγ*. The second terms in these expressions account for the influence of the edge cells on the second layer cells. The respective sums are over all neighboring edge cells that are moving inwards. Parameter *γ* is the angle formed by the direction of the line connecting the second-layer cell and the neighboring edge cell moving toward it, and *β* is the angle between the neighbor cell’s polarization direction and direction *γ*. Parameters *F*_*i*_ take the value of *F*_*edge*_.

## RESULTS

### EF guides both individual cells and groups of cells to the cathode

Typical physiological range of EF in wounds and embryonic tissues is 0.1-10 V/cm [41]. We investigated keratocyte groups moving in EF from this range, of magnitude ∼ 1 V/cm, and measured displacements of the groups of various sizes, as well as of single cells for up to 2 hours (Fig. 1A). We observed that at this time scale, the order of cells within the groups remained fixed – individual cells did not exchange neighbors, thus the cells in the groups were cohesive with their neighbors. Yet, the groups were not rigid, their shape deformed (Fig. 1A). Thus, the groups’ rheology combines fluid and solid features, in line with many previous observations [42]. Both single cells and groups of all sizes migrated to the cathode, which is clear from comparing earlier and later positions of groups of all sizes and shapes (Fig. 1A) and from trajectories of the groups’ centroids (Fig. 1B).

**FIGURE 1:**
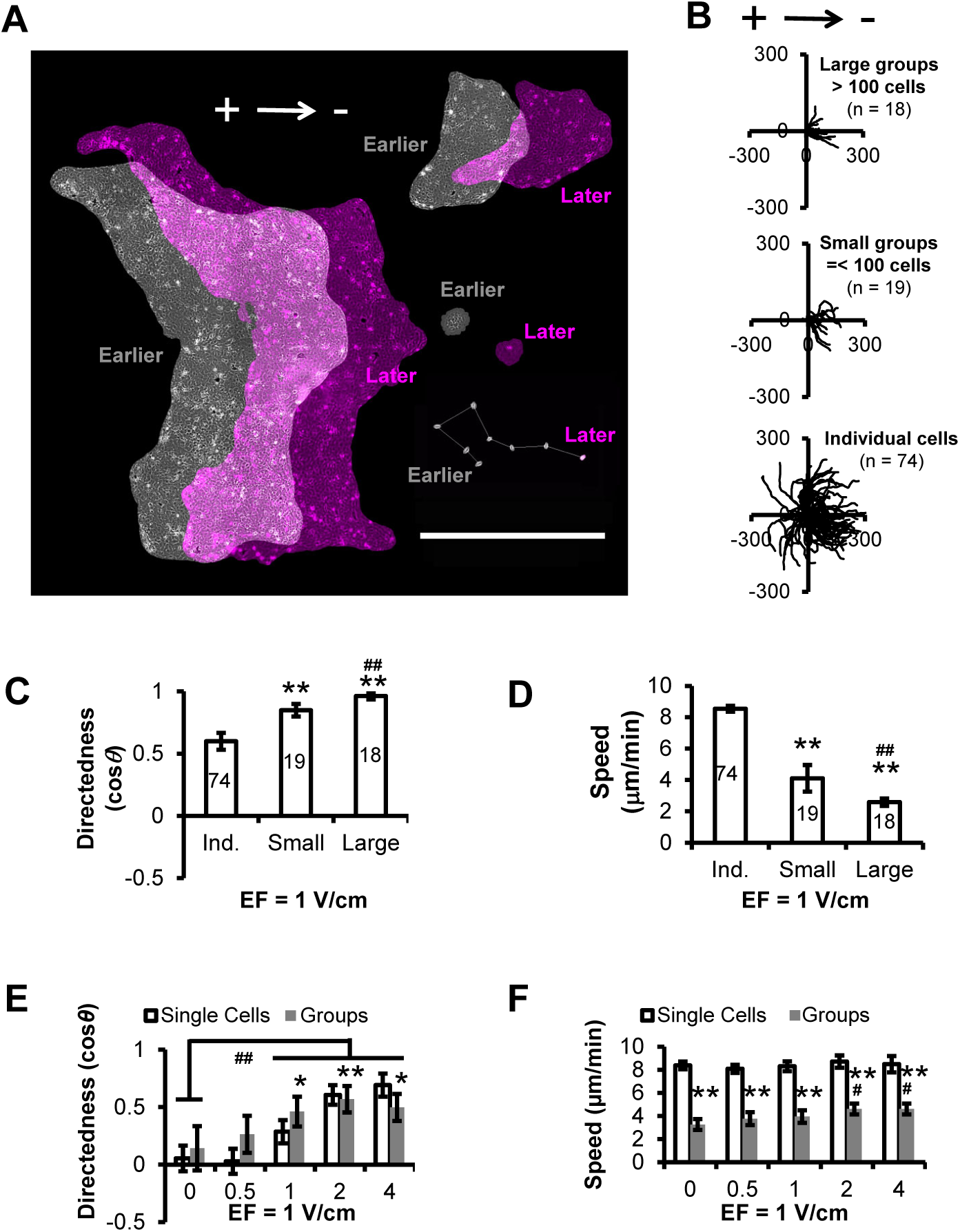
EF guides both single cells and cell groups to the cathode. A. Overlays of representative large and small cell groups and of one individual cell (with its migratory trajectory) before (gray) and 2 hrs after (magenta) applying 1 V/cm EF. Scale bar, 1 mm. B. Migration trajectories of large (more than 100 cells), small (less than 100 cells) groups and individual keratocytes after applying 1 V/cm EF for 30 minutes (cathode is at the right). The distances along x and y axes are in μm. C, D. Directedness (cos*θ*, see Methods) and speed (in µm/min) of individual cells, small and large groups, trajectories of which are shown in B. Data is presented as mean ± S.E. Numbers of quantified trajectories are indicated inside each bar. *\*\**, p < 0.01 compared to individual cells by unpaired Student t-test. ##, p < 0.01 compared to small groups by unpaired Student t-test. E, F. Directedness and speed of individual cells under different EF strength. Data is presented as pooled mean ± S.E. of at least three experiments. *\**, p < 0.05, combined groups compared to individual cells by unpaired Student t-test. *\*\**, p < 0.01 combined groups compared to individual cells by unpaired Student t-test. ##, p < 0.01 compared to No EF control by unpaired Student t-test.

A universal feature of cell groups migrating in EF or in chemical gradients is that directionality of the groups improves with size – larger groups move more persistently and faithfully along the external cue [9, 17, 18, 21]. We use directedness, cosine of the angle between the vector connecting the initial and final positions of the groups’ centroids and the cathode direction, as the measure of the directionality. One simple explanation is that the cells in the group tend to align with each other, which effectively filters out a directional noise [3, 43-45]. Similarly, we found that the groups are more directional than single cells, and that directionality improves with the group size (Fig. 1C).

Group speed, on the other hand, is not a universal function of the group size. Some studies reported faster migration of greater-size groups [9], others – that the group’s speed is largely independent of its size [46], yet others – that single cells move faster than groups [26]. Theoretical models can explain, as the experimental measurements indicate, that both increase and decrease of group’s velocity with its size are possible [45, 47]. In this study, we found that single cells moved faster than the groups, and the groups’ speed decreased with their size (Fig. 1D). Many factors could explain this effect: groups could move as slow as the slowest cell in the group, and/or equalizing speeds of individual cells in cohesive groups could involve interactions slowing down the movements.

Integration of individual cells into cohesive groups can lead to the groups higher sensitivity to directional signals than single cells [47]. For example, it was reported in [17] that isolated cells did not detect a weak EF, but cell groups did. We found that indeed the groups are more sensitive to EF than single cells (Fig. 1E) at EF of 0.5 and 1 V/cm. However, the sensitivity to EF becomes similar for the groups and individual cells at EF of 2 and 4 V/cm. The thresholds of sensitivity, between 0.5 and 1 V/cm for the groups, and between 1 and 2 V/cm for single cells, are similar. Considering that there are tens to hundreds of cells in each cohesive group, it is unlikely that a nontrivial synergy or a different EF sensing mechanism exist in the groups compared to single cells. Similar results were reported in [18, 23]. Speed of both single cells and cell groups was largely independent of the EF magnitude (Fig. 1F). Thus, the limiting factor for speed of migration is still the intrinsic cytoskeletal machinery of the cell. The presence of EF is but a polarization and directional cue.

### EF guides single PI3K-inhibited cells to the anode but groups of cells – to the cathode

As reported before [24, 25], after inhibition of PI3K by LY294002 compound, single cells migrate to the anode (Fig. 2A, B, C and E). We found that small (less than 100 cells) PI3K-inhibited groups are not directional, or more precisely, some of them shifted to the cathode, others – to the anode (Fig. 2A, B, C and E). Speed of these small groups was significantly smaller than speed of the unperturbed groups of similar size in EF (compare Fig. 1D and Fig. 2D). Large (more than 100 cells) PI3K-inhibited groups migrated to the cathode (Fig. 2A, B, C and E), same as large unperturbed groups. Speeds of single cells and of large groups after the PI3K inhibition were like those of unperturbed cells and large groups, respectively (compare Fig. 1D and 2D). Directedness of large PI3K-inhibited and unperturbed groups was similar (compare Fig. 1C and 2C).

**FIGURE 2:**
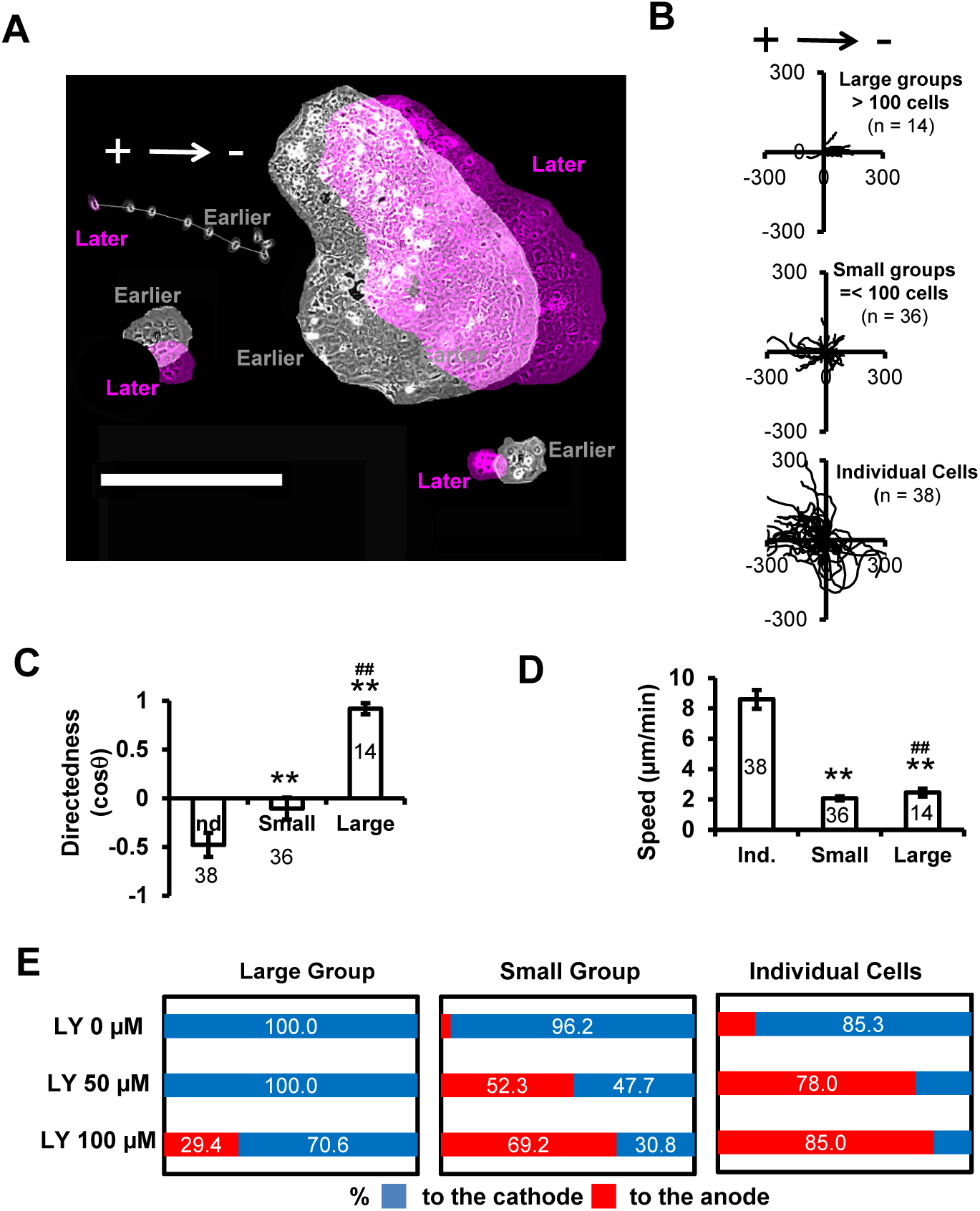
Effects of PI3 Kinase inhibition on cell group migration. A. Overlays of representative large and small cell groups and of one individual cell (with its migratory trajectory) before (gray) and 2 hrs after (magenta) applying 1 V/cm EF in the presence of 50 µM of LY292004 compound. Scale bar, 500 µm. B.Migration trajectories of large (more than 100 cells), small (less than 100 cells) groups and individual keratocytes after applying 1 V/cm EF for 30 minutes (cathode is at the right) in the presence of 50 µM of LY292004 compound. The distances along x and y axes are in μm. C, D. Directedness (cos*θ*, see Methods) and speed (in µm/min) of individual cells, small and large groups, trajectories of which are shown in B. Data is presented as mean ± S.E. Numbers of quantified trajectories are indicated inside each bar. *\*\**, p < 0.01 compared to individual cells by unpaired Student t-test. ##, p < 0.01 compared to small groups by unpaired Student t-test. E. Summary (percentage bar charts) of the displacement directions for individual cells, small and large groups after applying 1 V/cm EF for 30 minutes in the absence and presence of 50 and 100 µM of LY292004 compound.

Inhibition of myosin was very informative for deciphering rules of galvanotaxis for single keratocyte cells [25], so we used the drug blebbistatin to inhibit myosin contraction in cell groups. However, upon inhibition of myosin, cohesion between the individual cells in the groups was lost (Fig. S3A), likely because myosin contraction is necessary for stabilizing cell-cell adhesions [48]. Similar result was reported in [26]. Treatment of cell groups with the drug EGTA, which inhibits cell-cell adhesions, also led to disintegration of the group (Fig. S3B).

### Possible explanations of collective galvanotactic directionality

Obviously, these results cannot be explained if each cell in the group senses EF as a single cell of the same chemical state. Indeed, in this case, all cells of a PI3K-inhibited group would be polarized to the anode, and so the whole group would move to the anode, contrary to the data. To explain these results, it is informative to consider two simple conceptual models of collective directional cell motility [47]. Two more models are considered and argued against in the Discussion; the list of models we consider is by no means complete.

One possibility is that only cells at the edge of the group (cells that have a free cell edge that is not in touch with edges of other cells, hereafter called simply ‘edge cells’) are sensing the directional cue; cells inside the group (cells surrounded on all sides by other cells, from now on called ‘inner cells’) follow the collective action of the edge cells passively (Model 1, Fig. 3A). More specifically, one possible assumption would be that the edge cells’ polarization is determined only by lamellipodial edges of these cells at the group’s boundary, and that the lamellipodia are always directed to the cathode, whether in uninhibited, or in PI3K-inhibited cells. (Another part of this assumption would be that it is the cell body that directs the cell to the anode in single cells, and this effect re-directs single PI3K-inhibited cells to the anode, but the cell body effect is completely inhibited by mechanical contacts with other cells.) It would be also natural to assume that this bias to the cathode of the edge cells is mediated by the geometry of the group’s boundary: edge cells at the cathodal side of the group, for which lamellipodia are facing the cathode, are polarized the most, while edge cells at the anodal side of the group, for which lamellipodia are facing the anode, opposite to where the lamellipodia ‘would want to go’, are polarized the least. This model then suggests the distribution of the motile ‘drives’ for the edge cells depicted in Fig. 3A. This model predicts (see the detailed model description in the Methods and in the next section), correctly, that both unperturbed and PI3K-inhibited groups would move to the cathode (Fig. 3A). This model also makes two other nontrivial predictions, true in both unperturbed and PI3K-inhibited cases: a) Small groups with similar numbers of inner and edge cells should move to cathode, whether unperturbed, or PI3K inhibited, which is contrary for the data for the PI3K-inhibited groups. b) Computational model predicts that in this case the velocity distribution is such that the cell speed inside the group increases along the anode-cathode axis and reaches a maximum at the frontal, cathodal side of the group (Fig. 3D).

**FIGURE 3:**
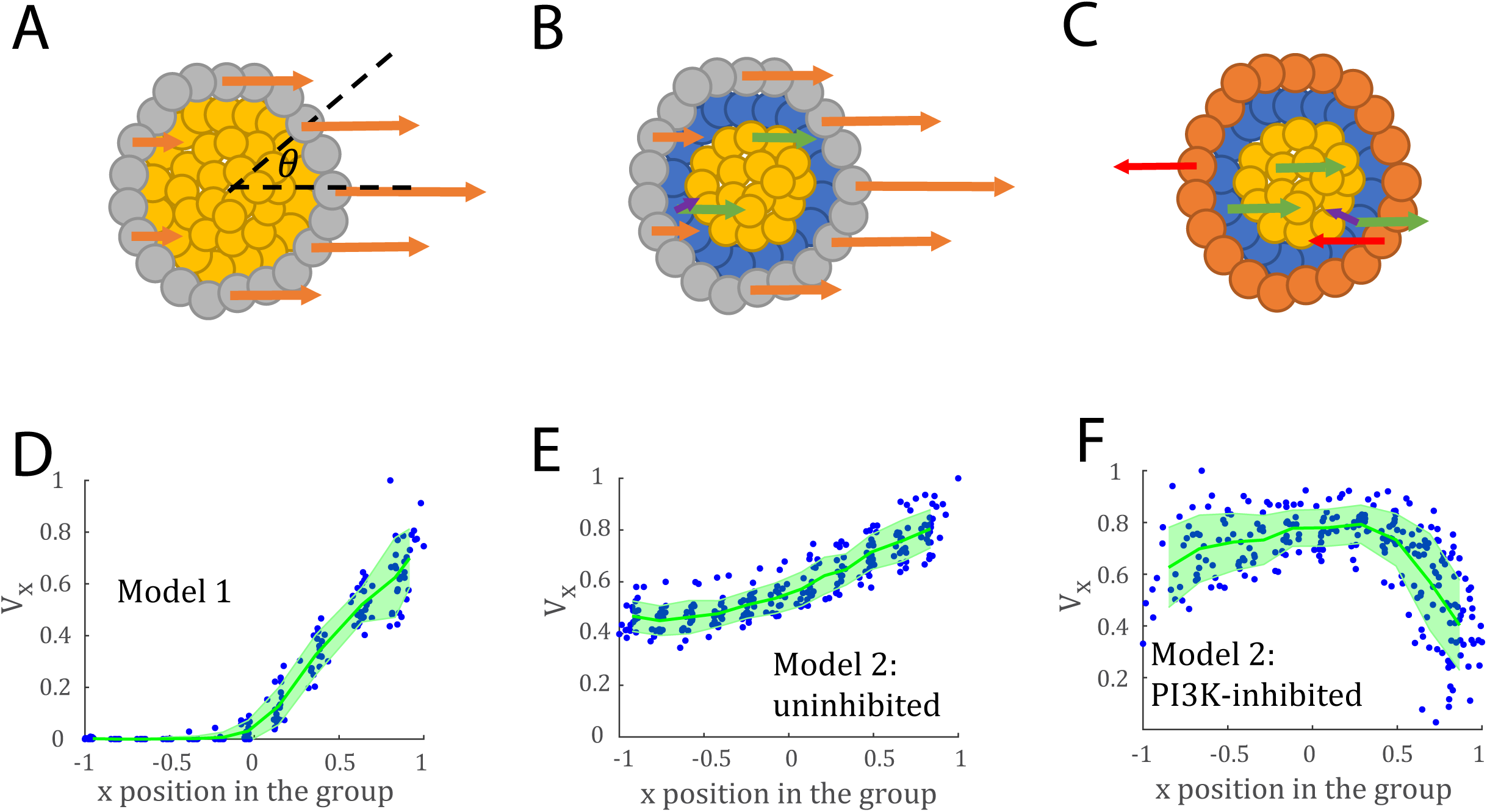
Two models and cell predicted velocity distribution within groups. A-C. The schematic diagrams showing the cell driving forces for two models: ‘only-edge-cells-active’ model 1 (A) and ‘both-inner-and-edge-cells-active’ model 2 (B, C). Cathode is always on the right. In (A) (for both unperturbed and PI3K-inhibited cells), the edge cells (grey) sense EF leading them to the cathode, and the driving forces (orange arrows) at the front of the group are larger than those at the back. The inner cells (yellow) do not have active forces and are pulled or pushed by the edge cells. In (B) (for unperturbed cells), the edge cells are the same as in (A) while the inner cells also have driving forces pointing to the cathode (green arrows). In addition, the second-layer inner cells (the blue cells contacting the edge cells directly) are influenced by the edge cells that attempt to move inward and have another force component (the purple arrow). In (C) (for the PI3K-inhibted cells), the forces of the edge cells point to the anode, while the inner cells are the same as in (B). Similarly, the second-layer cells are also influenced by the neighboring edge cells. D-F. Distributions of x-components of cell velocities along the anode-cathode diameter for the ‘only-edge-cell-active’ model 1 (D, corresponding to model 1, A), and the ‘both-inner-and-edge-cells-active’ model for unperturbed cells (E, corresponding to model 2, B) and for PI3K-inhibited cells (F, corresponding to model 2, C). The data to calculate this distribution is obtained from 10 simulations using cells within the rectangular box shown in Fig. S2 within 200-computational time units-long window. Each blue dot is a data sample and the green line is the average of the data samples within the same grid as that shown in Fig. S2. The shaded areas show the standard deviations. The distance across the group is normalized so that -1 and +1 correspond to the rear and front, respectively. The velocity is normalized so that 1 is the maximal velocity in the group.

Another, more complex model, is that inner cells sense EF, and always polarize to cathode, whether unperturbed or PI3K-inhibited. Edge cells behave roughly as single cells and therefore pull to the cathode, in sync with the inner cells in unperturbed groups (Model 2, Fig. 3B) but push to the anode in PI3K-inhibited groups (Model 2, Fig. 3C). This model predicts, correctly, that both unperturbed and PI3K-inhibited *large* groups move to the cathode, because in such groups inner cells are in majority and win in the directional tug-of-war. This model also makes two other predictions: a) Small groups, with similar numbers of inner and edge cells, move to the cathode in the unperturbed case, when both inner and edge cells are driven to the cathode, but are not directional in the PI3K-inhibited case, because inner and edge cells pull and push in opposite directions and roughly balance each other mechanically. b) Speed of individual cells in unperturbed groups is distributed in space like the prediction of the previous model (Fig. 3E), but in PI3K-inhibited groups, cell speeds should be higher at the group’s center, because the cells at the front act as breaks on inner, cathode-directed, cells.

To see if the qualitative predictions of these two models are also valid quantitatively, we used Cellular Potts Model (CPM, see Methods for details), which simulates cells as deformable bodies, adhering to each other [49-51]. The shapes and movements of the group cells are governed by an effective balance between intracellular pressure, cortex tension and cell-cell adhesion. Effectively, the model mimics complex viscoelastic interactions between neighboring cells [49-51]. To the standard CPM, we added effective elastic coupling between neighboring cells and an effective force driving individual cells to or away from the cathode. Similar motile force was used in CPM before [44, 52, 53].

Simulations of ‘only-edge-cells-active’ Model 1 showed that in this case, as expected, speed at the group front is maximal (Fig. 3D). Note that this simulation covers both unperturbed and PI3K-inhibited groups. Simulations of ‘both-inner-and-edge-cells-active’ Model 2 for the unperturbed group predicted that in this model the speed profile is also characterized by the maximal cell speed at the group front (Fig. 3E), providing the assumption that the edge cells at the cathodal edge of the group pull stronger than the inner cells. Finally, simulations of this Model 2 for the PI3K-inhibited group shows, as intuited, that in this case cells in the middle of the groups are the fastest (Fig. 3F).

### Unperturbed groups are pulled from the front; PI3K-inhibited groups are pushed from the center

PIV analysis showed that in large unperturbed groups, 10 to 30 minutes after the EF was switched on, speed of the cells at the front of the groups was highest (Fig. 4A, C and E). The maximal cell speed in large PI3K-inhibited groups was observed at the rear and center of the groups (Fig. 4B, D and E), in agreement with the ‘both-inner-and-edge-cells-active’ model. We also observed that the cells in unperturbed groups respond to the EF directionally synchronously in all parts of the group (Fig. S4D), which is in line with this model as well.

**FIGURE 4:**
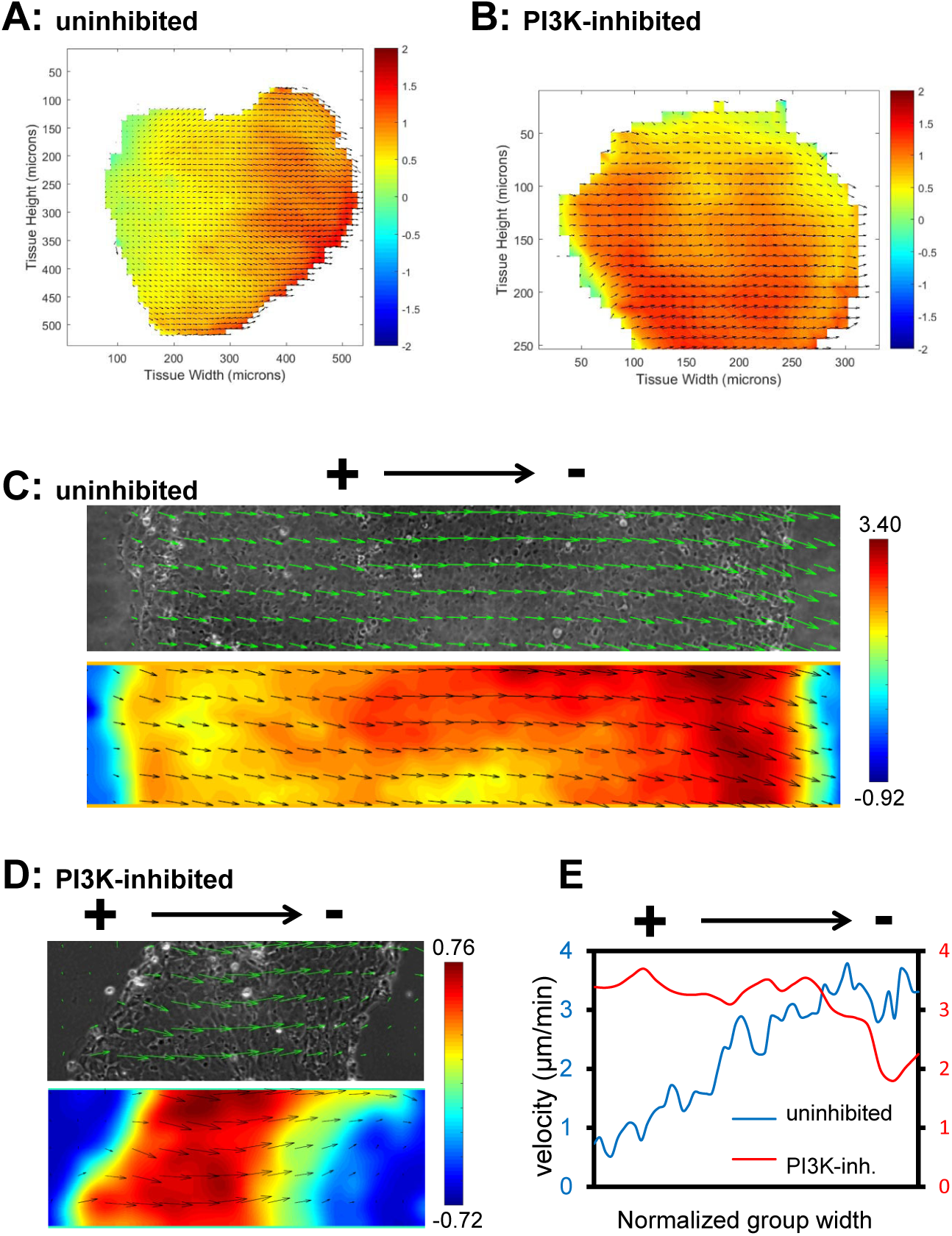
Cell Velocity distributions inside the migrating cell groups. A, B. Spatial velocity maps of large uninhibited (A) and PI3K-inhibited (B) cell groups undergoing EF-directed migration to the cathode. Component of the velocity vector parallel to the cathode direction (to the right) was averaged in time in the interval between 20 and 30 minutes after application of 1 V/cm EF. The averaged EF-directed component of the velocity (in µm/min) spatial distribution is color-coded; also, the velocity vectors measured 30 minutes after application of 1 V/cm EF are shown. C, D. Same as (A, B), but the velocity (in µm/min) distributions are shown within ∼ 100 μm-wide slices of the central parts of migrating cell groups. The cathode is at the right; the rears and fronts of the groups are at the left and right, respectively. E. Line plots of the EF-directed components of the velocities shown in C and D. The EF-directed components of the velocities were integrated across the widths (in the direction normal to EF) of the groups’ slices shown in C and D; the distances along the lengths of these slices were rescaled so that left and right ends of the graph correspond to the rear and front of the groups, respectively.

In the PI3K-inhibited groups, cells at the center always tended to move to the cathode, while at the periphery and in long narrow appendices of the groups cells transiently tended to move to the anode, oppositely to the whole group (Fig. S4A-C). In fact, in several cases we observed that a small sub-group from the edge of the PI3K-inhibited group even split off from the main group and was left behind (Fig. S4E). This, again, argues in favor of the model according to which edge cells behave actively and directionally, like single cells.

### Computational model accounts for all experimental observations

To see if the computational model could account for the data quantitatively, we made the motility of the cell noisy and searched for mechanical and directional parameters for inner and edge cells that generated realistic trajectories of individual cells. We then simulated movements of the single cells and small and large groups in both unperturbed case (Fig. 5) and PI3K-inhibited case (Fig. 6) for Model 2. We found that the calibrated model accounts for all our experimental observations and measurements quantitatively. Namely, the model mimics realistic group displacements (compare Fig. 5A and 6A with Fig. 1A and 2A, respectively). The model also generates realistically looking trajectories of the single cells and small and large groups (compare Fig. 5B and 6B with Fig. 1B and 2B, respectively). Lastly, the model correctly predicts the directedness and speed of the groups as well (compare Fig. 5C-D and 6C-D with Fig. 1C-D and 2C-D, respectively).

**FIGURE 5:**
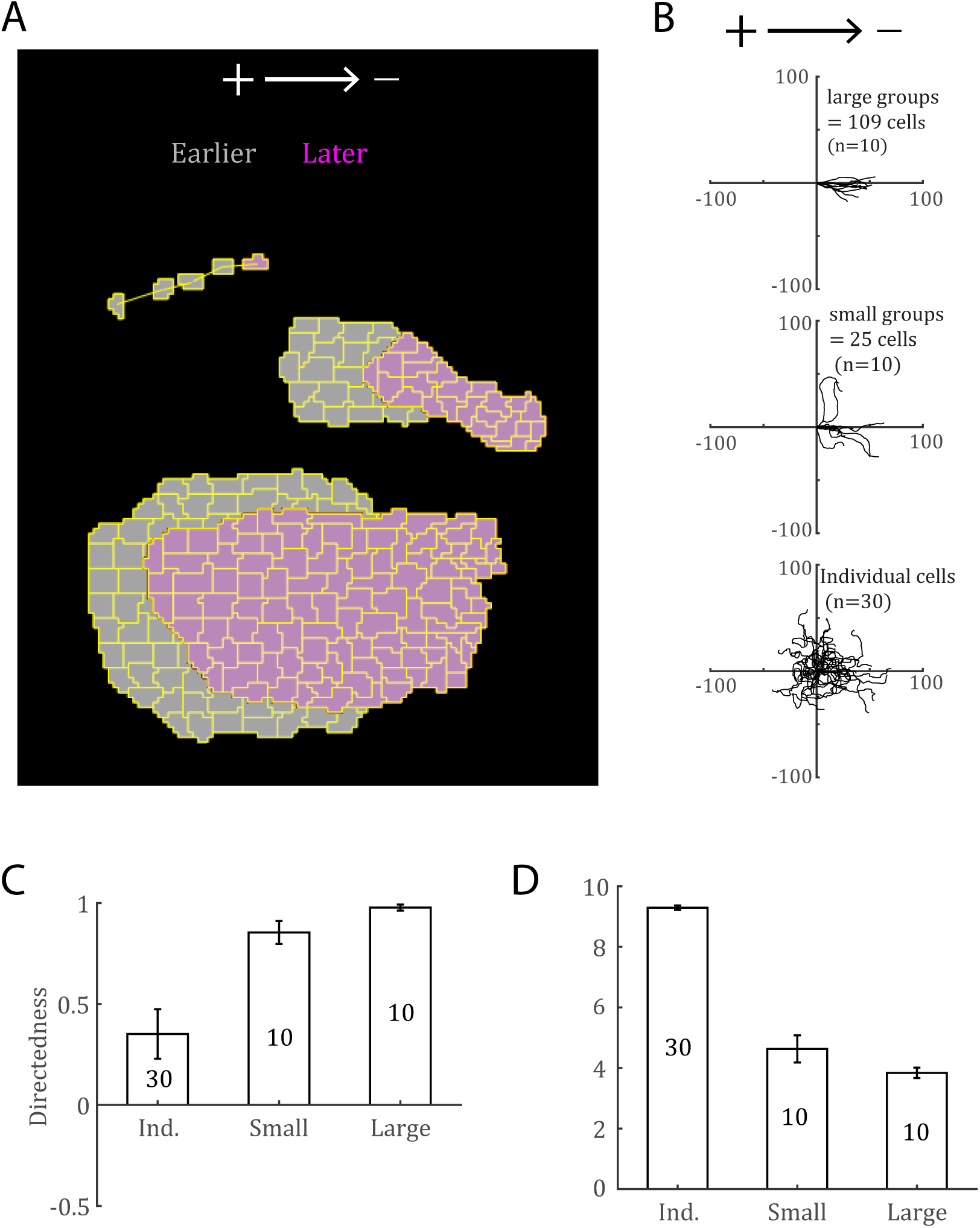
EF guides both single cells and cell groups to the cathode in simulations, corresponding to the experimental results in Fig. 1. A.Overlays of representative large and small cell groups and of one individual cell (with its migratory trajectory) earlier (gray) and later (magenta). The snapshots are taken at times 50 and 250 for the cell groups and at times 50, 100, 150, 200, 250 for the individual cell. All times are in units of computational time step. B.Migration trajectories of large (109 cells) and small (25 cells) groups and individual cells. There are 10 trajectories for the cell groups and 30 trajectories for the individual cells. All the trajectories and the following calculation in C and D are using data from time 500 to 999. C, D. Directedness (*cosθ*) and speed of individual, small and large groups, trajectories of which are shown in B. The error bar shows the standard error. The length unit in the simulation is pixel and the time unit in the simulation is Monte-Carlo Step. We normalize the speed to make the value comparable to that in the experiment.

**FIGURE 6:**
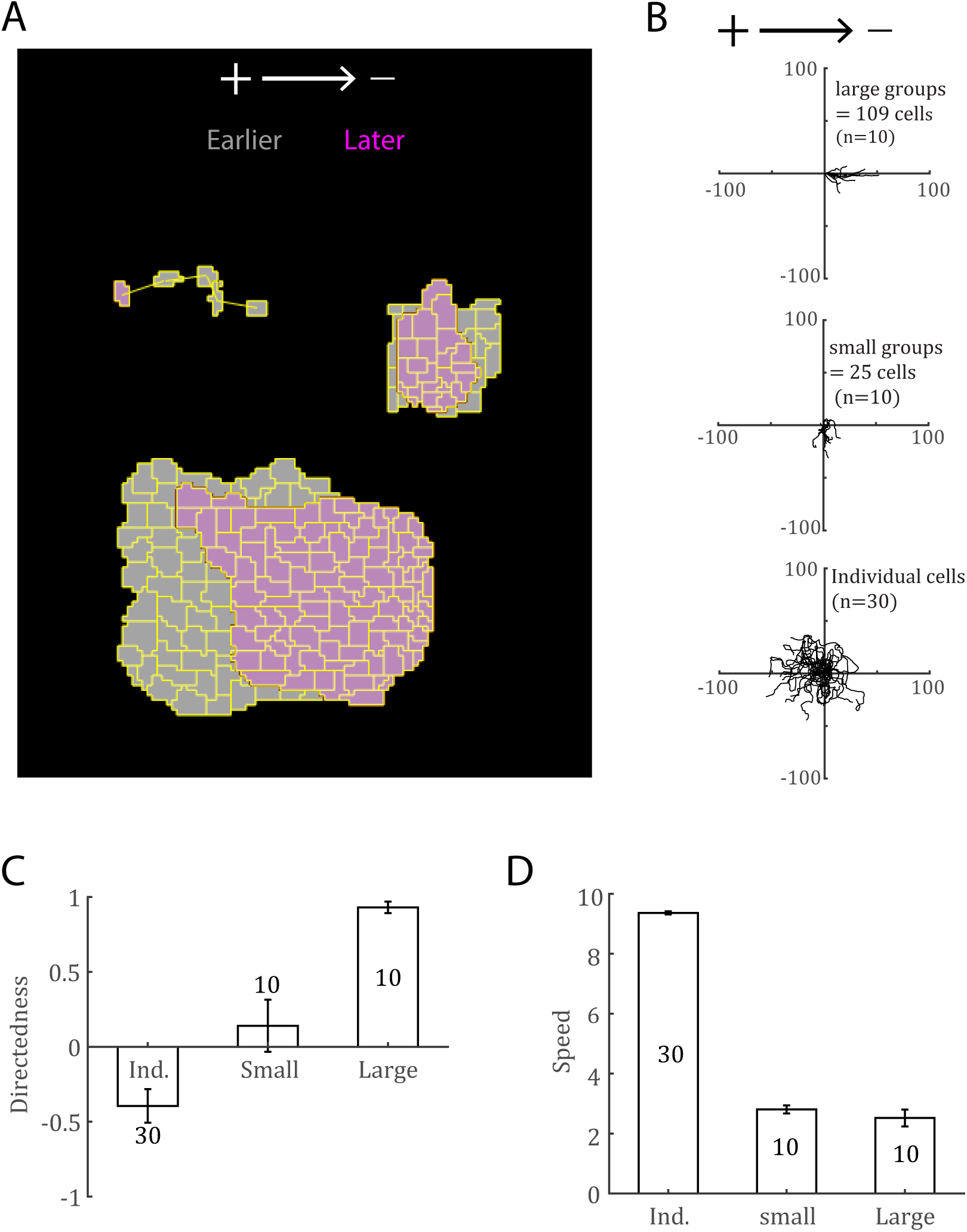
Effects of PI3K on cell group migration in the simulation, corresponding to the experimental results in Fig. 2. A.Overlays of representative large and small cell groups and of one individual cell (with its migratory trajectory) earlier (gray) and later (magenta). The snapshots are taken at time 50 and time 250 for the cell groups and at time 50, 100, 150, 200, 250 for the individual cell. All times are in units of computational time step. B.Migration trajectories of large (109 cells), small (25 cells) groups and individual cells. There are 10 trajectories for the cell groups and 30 trajectories for the individual cells. All the trajectories and the following calculation in C and D are using data from time 500 to 999. C,D. Directedness (*cosθ*) and speed of individual, small and large groups, trajectories of which are shown in B. The error bar shows the standard error. The length unit in the simulation is pixel and the time unit in the simulation is Monte-Carlo Step. We normalize the speed to make the value comparable to that in the experiment.

These results can be qualitatively understood as follows: directedness increases with the group size because of, first, the effective averaging effect caused by the mechanical adhesion between neighboring cells, which filters out the noise in the individual cell’s directions; and second, the alignment effect between neighboring cells. Both effects are greater for larger groups due to the law of large numbers. On the other hand, the group speed decreases as the group size increases, because the mechanical adhesion between the neighboring cells that try to move with inherently different speeds and directions, effectively slow down the interacting cells while preventing their detachment. The larger the group, the more this effect is pronounced.

Finally, we simulated a group with a narrow finger-like appendix in the PI3K-inhibited case (Fig. S4F). Indeed, in the simulation, this appendix was left behind the greater and rounder part of the group and eventually detached from it (Fig. S4F), much like observed (Fig. S4E).

## DISCUSSION

### Summary of the experimental results

The question about mechanisms of the collective cell direction sensing has led us to investigate how cohesive keratocyte groups of various sizes move in EF. We found that the groups’ sensitivity to weak EF is only slightly better than that of the single cells. Expectedly, single unperturbed cells and groups of all sizes crawl to the cathode. Group’s speed decreases with its size, while its directionality improves with the size. Our main finding is that large groups of PI3K-inhibited cells move to the cathode, oppositely to single PI3K-inhibited cells that move to the anode. Though both large uninhibited and PI3K-inhibited groups move to the cathode, the cell speed distribution in these groups is different: cell speed reaches maximum at the front of the uninhibited groups, but at the front of the PI3K-inhibited groups cell speed is minimal. Small groups of PI3K-inhibited cells do not have a preferred direction of migration. We observe that in PI3K-inhibited groups, edge cells and cells in long parts of the groups have a transient tendency to move to the anode.

### Theoretical model explaining the data

The simplest explanation of the data is that inner cells always, independently of the cells’ chemical state, tend to go to the cathode, while the edge cells behave directionally like single cells. Indeed, in large PI3K-inhibited groups with complex shapes, small cell clusters at the groups’ periphery sometimes show tendency to go to the anode, oppositely to the cathodal direction of the main part of the group. According to this model, unperturbed cells groups move to the cathode, as all cells in the group are driven to cathode, and the cell speed is maximal at the front because the edge cells at the front are polarized to the cathode stronger than the inner cells. On the contrary, the directionality of PI3K-inhibited cell groups is determined by a mechanical tug-of-war between the cathode-driven inner and anode-biased edge cells. In larger groups, the former win and the group migrates to cathode. At the front of such groups, the edge cells are weakly polarized to the anode, against the majority of the inner cells pushing to the cathode, and so the cell speed is higher in the middle and lower at the front of the groups. In small PI3K-inhibited groups, the oppositely polarized inner and edge cells are roughly balances, and the groups are not directional.

### Can single cells and groups move oppositely in response to other directional cues?

It would be interesting if the phenomenon we discovered – that under some conditions individual cells and cell groups migrate in opposite directions in response to a directional cue – is not unique for the galavanotaxis, and that there is an analogous phenomenon in, for example, chemotaxis. We are not aware of exactly the same phenomenon in chemotaxis; the closest analogy is with one recent study [46] that reported that lymphocyte clusters always followed chemical gradient, independent of its steepness, while single cells reversed directionality for very steep gradients. It is well known in chemotaxis that individual cells can be turned around by some chemical perturbations [54], but how cell groups will behave under the same perturbations is unclear.

### Relation of our data to other studies

Our results have interesting parallels with data from two recent studies: Cohen et al. observed that cells at the boundary of epithelial cell groups migrating in EF did not respond to EF directionally [9] implying that the inner cells guide the whole groups. Similar observations were made for clusters of malignant cancer cells responding to chemical gradients [6]. Also related is the study [55] reporting that PI3K is upregulated in leader cells, and that when PI3K is inhibited, collective migration of MDCK epithelial cells is disrupted. Thus, it is possible that when PI3K is inhibited, this affects the edge cells with large lamellipodia the most, and in this case the inner cells take over the role of responding to EF making the whole group directional.

### Two other conceptual models are unlikely to explain the data

These two studies raise the question: is it possible that only inner cells, which are directed to the cathode in any chemical state, are leading the group, while the edge cells are following passively? Part of our data could be explain by such model, however, contrived assumptions would be needed to explain why the velocity profiles of the uninhibited and PI3K-inhibited groups are different, and why some edge cells in the PI3K-inhibited groups tend to transiently move to the anode. Thus, we argue that in our experimental system both inner and edge cells are active directionally, albeit in different ways.

Another possibility is that the cells inside the group neither move, nor sense EF individually, but are rather mechanochemically integrated into a large supracellular structure. Integrated cytoskeletal global networks in collectively migrating keratocytes [26] and in neural crest cell groups [11] were reported. Traction force measurements showed integrated changes in patterns of intercellular stresses that precede changes of individual cell shapes [21] in the groups in EF indicating supracellular response. Such integration could effectively enable the group to sense EF globally, on the length scale of the whole group. Indeed, cells can read the bioelectrical state of distant regions in the group via the chemical molecules redistributed across long distances by a gradient of bioelectric cell state [56]. There is also a possibility of global measurement of group size and of global coordination through mechanical stresses or diffusing morphogens [47].

This model is hard to rule out definitively, but we would like to offer the following argument against it. As noted in the Introduction, electrophoresis generates a gradient of charged mobile transmembrane proteins across the keratocyte cell in EF [24]; it is likely that the cell measures this gradient and converts it into the directional motile behavior [24]. If the cell size is equal to *l*, EF strength is equal to *E* and effective charge of the mobile protein is equal to *q*, then the ratio of the protein concentrations between the cathodal and anodal ends of the cell is on the order of *exp* (*qEl* / *k*_*B*_*T*) [24], where *k*_*B*_*T* is the thermal energy (*k*_*B*_ is the Boltzmann’s constant, and *T* is the absolute temperature). If the cells in the groups are integrated into the supracellular structure, they could in principle measure the gradient of the mobile charged proteins across the whole group. If *L* is the group’s size (for round group, 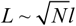 where *N* is the number of cells in the group), then the ratio of the protein concentrations between the cathodal and anodal ends of the group is on the order of *exp* (*qEL* / *k*_*B*_*T)*. If *R* is the weakest protein front/rear concentration ratio that could be detected, and it is similar for the single cell and the super-cell, then the threshold EF that could be detected is inversely proportional to the size of the cell and super-cell, respectively. Indeed: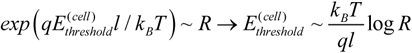. As cell groups are an order of magnitude greater in size than single cells, then according to this argument, the super-cell could sense EF, which is at least an order of magnitude weaker compared to that sensed by individual cells:

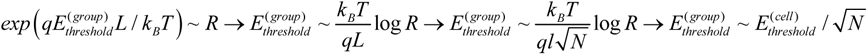

(recall that 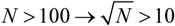).However, this is not the case (Fig. 1E).

### Important question that remains open is: why inner and edge cells interpret EF differently?

There are significant differences in cytoskeletal organizations in inner and edge cells [26], the most prominent of which is large lamellipodia of the edge cells. Usually, inner cells have only cryptic lamellipodia [57]; active lamellipodia of these cells are suppressed by physical coupling to neighboring cells on all sides [21]. Intercellular junctions and cadherin and planar cell polarity signaling [17, 58] also have different geometric organizations in inner and edge cells. Due to these different organizations of cytoskeletal and signaling networks inside the group and at its edge, it is possible that the balance of the signal transduction pathways [25, 59] in inner cells differs from that in single and edge cells.

In [25], we proposed a ‘compass’ model of galvanotaxis, according to which a strong, PI3K-dependent EF-sensing pathway polarizes motile cell to the cathode, while a weaker PI3K-independent pathway orients the cell to the anode. We suggest the following modification of this model that explains our results (Fig. 7).

**FIGURE 7:**
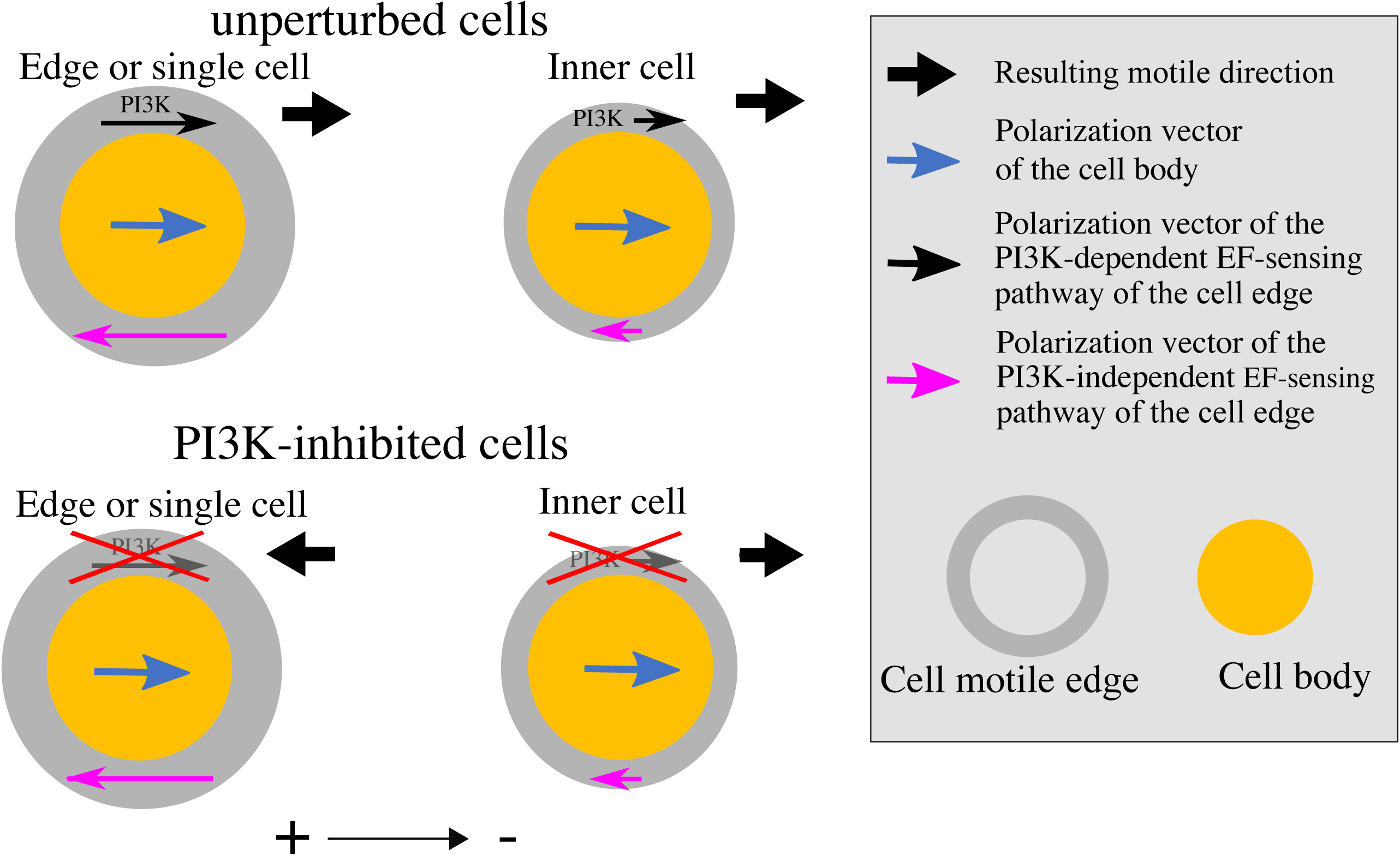
Hypothesized balances of the polarization signals in cells. Possible balance of three polarization vectors, two at the cell motile edge, and one in the cell body are shown. The hypothesis is that these vectors add geometrically to determine the resulting locomotory direction. The polarization vectors’ magnitudes depend on the cell position in the group (edge or inner cell) and on the cell chemical state (uninhibited or PI3K-inhibited). Note that the motile edge and respective polarization signal are larger and stronger, respectively, in the single or edge cell compared to that in the inner cell.

We propose that EF signal is transduced by three pathways – one to the cell body, and two other, a PI3K-dependent one and a PI3K-independent one, to the cell’s motile edges. Each of these pathways contributes to the cell directional polarization in an additive way. We characterize the strengths of these pathways by abstract ‘polarization vectors’, the length of which is proportional to relative weight of the respective pathway in determining the cell direction. *P*_*cb*_ is the cathode-directed polarization vector of the cell body; *P*_*me*_ is the anode-directed polarization vector of the PI3K-independent pathway to the motile edge; *P*_*mePI3K*_ is the cathode-directed polarization vector of the PI3K-dependent pathway to the motile edge. All we assume is that two pathways to the motile edge of the single cell are roughly balanced and are stronger than the pathway to the cell body: |*P*_*me*_| ≈ |*P*_*mePI3K*_| > |*P*_*cb*_|. Then, in the single or group-edge uninhibited cell, the resulting polarization vector, (*P*_*me*_ + *P*_*mePI3K*_ + *P*_*cb*_), is directed to the cathode (because *P*_*me*_ + *P*_*mePI3K*_ ≈ 0, and *P*_*cb*_ is cathode-directed). In the single or group-edge PI3K-inhibited cell, the resulting polarization vector, (*P*_*me*_ + *P*_*cb*_), is directed to the anode (because |*P*_*me*_| > |*P*_*cb*_|, and *P*_*me*_ is anode-directed). In the inner cell, however, the motile edge of the cell is narrow, and we assume that the respective pathways are weaker, and so the polarization vectors, *p*_*me*_ and *p*_*mePI3K*_, become smaller: |*p*_*me*_| ≈ |*p*_*mePI3K*_| < |*P*_*cb*_|. Then, in the inner cells, whether uninhibited, or PI3K-inhibited, the resulting polarization vectors, ([*p*_*me*_ + *p*_*mePI3K*_ + *P*_*cb*_] and [*p*_*me*_ + *P*_*cb*_], respectively) are directed to the cathode (because *P*_*cb*_ is cathode-directed and is dominant in both cases).

### Limitations of the study

There is much complexity of the collective galvanotactic response that our study did not address. There are likely slow (on the scale of hours) processes of adaptation of the group migration to EF, which is manifested in gradual slowing down of the migrating groups that we observed after 2 hours of exposure to EF. Similar gradual slowing down of collective keratocytes’ groups migrating spontaneously was reported in [26]. One possible mechanistic clue to respective process is observed gradual redistribution of global stresses in the groups of HaCaT cells migrating in EF [21]. Other sources of complexity are natural cell-to-cell variabilities in directional responses, nontrivial rheology of the cell clusters and stochastic effects in collective directional sensing [60]. Future research will shed light on impacts of these complexities on collective galvanotaxis.

## CONCLUSION

Individual fish keratocyte cells migrate to the cathode, while inhibition of PI3K reverses single cells to the anode. Chemically unperturbed cell groups of any size move to the cathode. Large groups of PI3K-inhibited cells move to the cathode. Small groups of PI3K-inhibited cells are not directional. The fastest cells are at the front of the uninhibited groups, but at the middle and rear of the PI3K-inhibited groups. These results are most consistent with the model according to which inner cells, whether unperturbed or PI3K-inhibited, always have tendency to move to the cathode, while peripheral cells in the group behave as single cells: they are directed to the cathode if uninhibited, but biased to the anode, if PI3K-inhibited. A mechanical tug-of-war between the inner and edge cells directs large groups with majority of the inner cells to the cathode.

## AUTHOR CONTRIBUTIONS

Yaohui Sun, M.Z. and A.M. conceived and directed the study; Yaohui Sun, H.Y., M.Z. and A.M. wrote the manuscript; Yaohui Sun, M.Z. and A.M. analyzed the data with help from H.Y., K.Z., Y.Z. and F.L.; Yaohui Sun did the experiments with help from Yuxin Sun, X.G. and B.R.; H.Y. did the modeling with help from C.C.

## ACKNOWLEDGMENTS

This work was supported by US Army Research Office grant W911NF-17-1-0417 to A.M., by Air Force Office of Scientific Research grant FA9550-16-1-0052 to M.Z. (Program PI: Wolfgang Losert), and by National Institutes of Health grant EY 019101 to M.Z.

## Figure legends

**SUPPLEMENTAL FIGURE 1:**
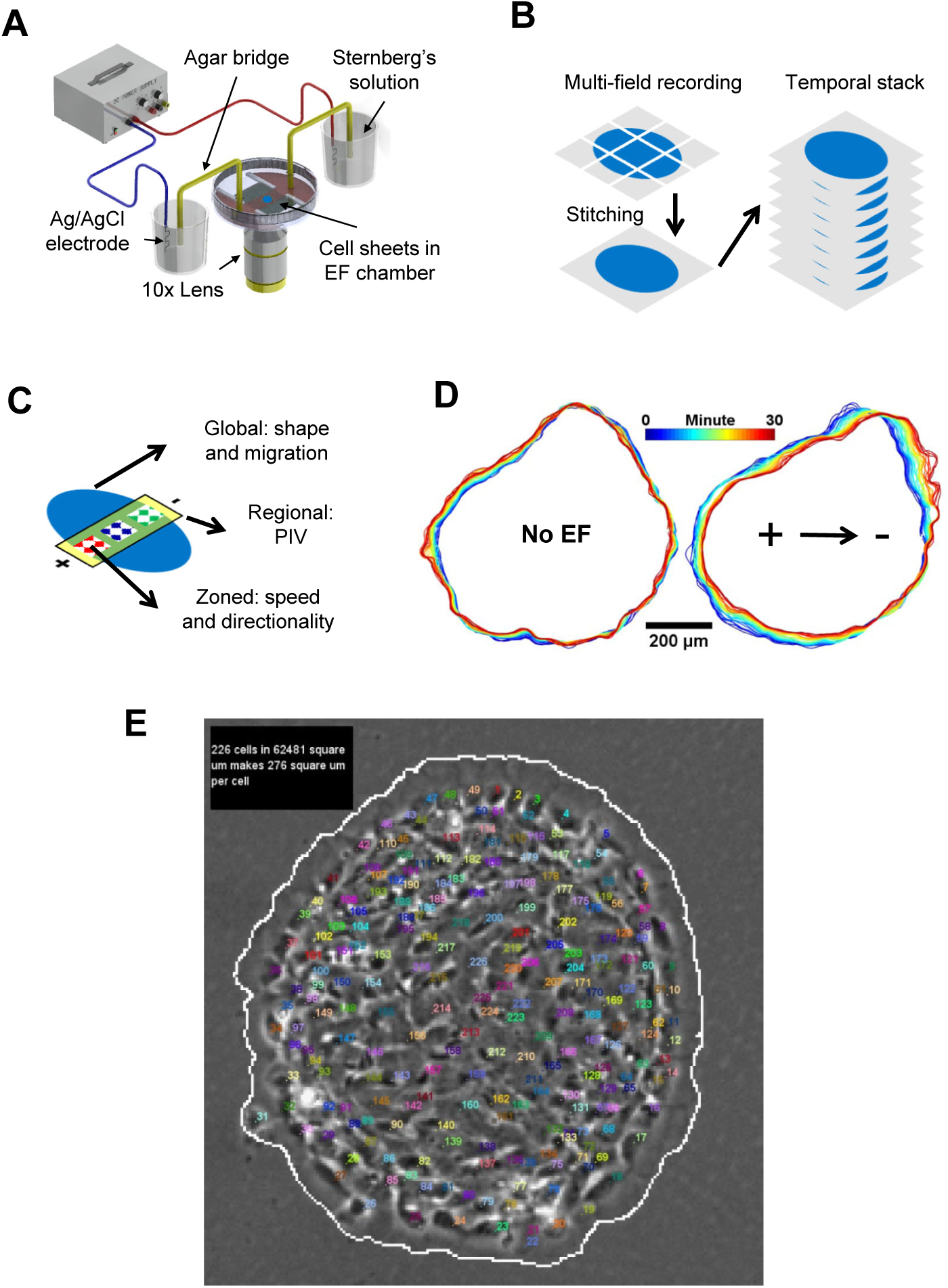
EF-guided collective cell migration of fish keratocytes. A. Schematic view of the electrotaxis chamber and the setup for EF application. B. Workflow for image processing. Cell group movement was monitored by overlapped multi-field time lapse recording. Multi-field images were stitched together as described in Methods and assembled into temporally ordered stacks for subsequent quantification. C. Schematic view of analytic approaches (global, regional and zonal analysis) for characterizing collective keratocyte migration. D. Cell group boundary contour overlays before and after EF application, each for 30 minutes in a five-minute interval. Contours are color coded as shown in the color bar. Scale bar, 200 μm. E. Example of enumerating and calculating cell number in the keratocyte group. Scale bar, 100 μm.

**SUPPLEMENTAL FIGURE 2:**
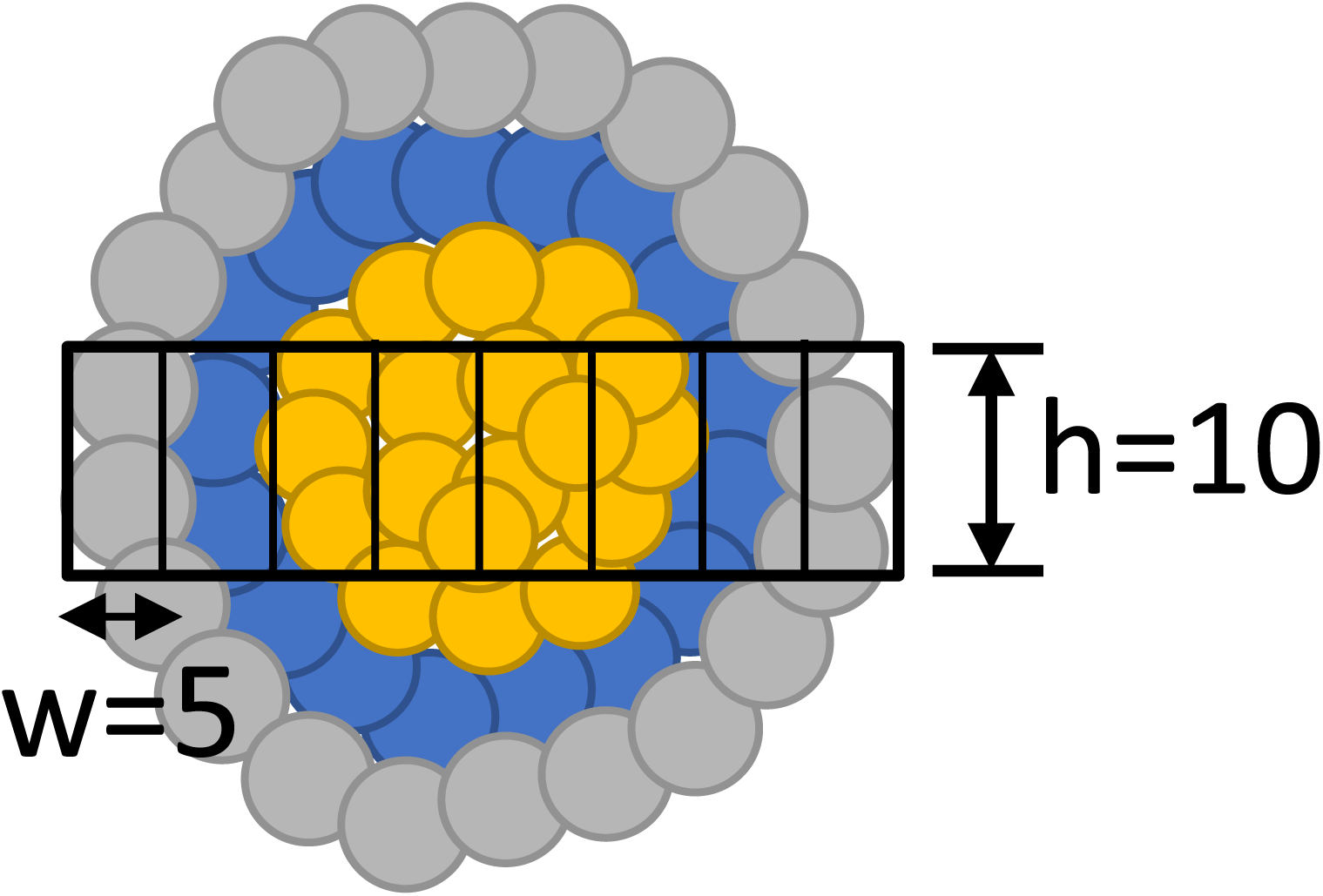
Calculating the velocity distribution shown in Fig. 3. The cells that are taken into account in the calculations are within the rectangular box along the central anode-cathode axis of the group of height 10 pixels. The whole box is divided into the grid with sub-box width 5 pixels. We take data from 10 simulations. For each simulation, for each cell within the rectangular box, we get its average x component of velocity as (*X*_*t*1_ − *X*_*t*0_)/(*t*1 − *t*0). We use *t*1 = 199 and *t*0 = 1 (time is measured in units of computational time steps) in the final results but the results will not change greatly as long as *t*1 is not too large, as after long enough time, the distribution of velocity within the group becomes uniform. We get the corresponding relative x-position using 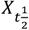 in which 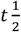 is in the middle of the *t*0 to *t*1 interval. All data samples of velocity and relative x position are averaged within each grid and shown as the green line in Fig. 3.

**SUPPLEMENTAL FIGURE 3:**
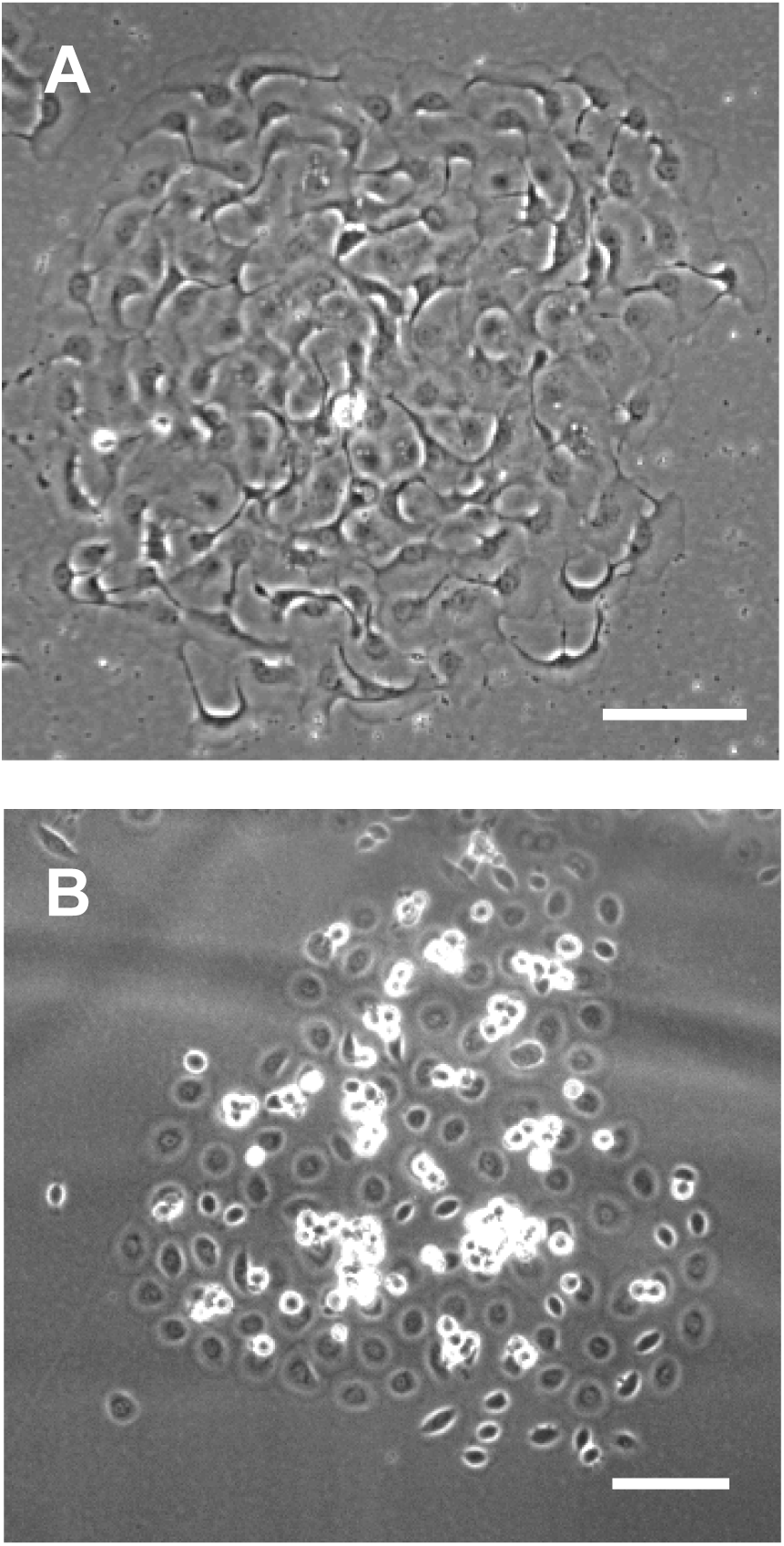
Blebbistatin and EGTA disrupt cohesion of the cell groups. A.Cohesion between individual cells in the keratocyte group is lost 15 min after application of 50 µM of Blebbistatin. Scale bar, 100 µm. B.Cohesion between individual cells in the keratocyte group is lost 15 min after application of 5 mM of EGTA. Scale bar, 100 µm.

**SUPPLEMENTAL FIGURE 4:**
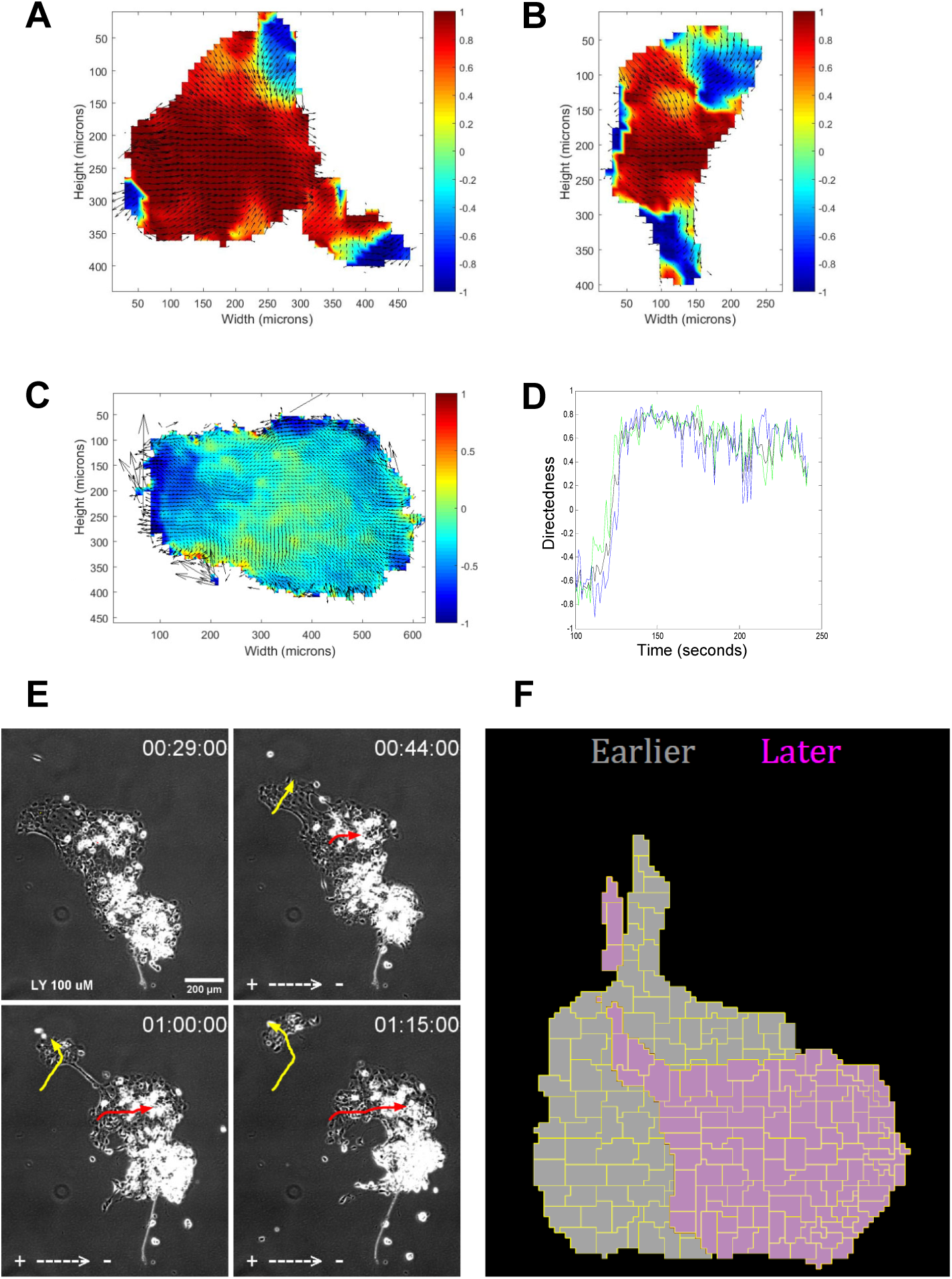
Regional behaviors of cell groups in EF. A - C. Spatial maps and velocity vectors of directedness measured at various moments within 30 min time interval for three different keratocyte groups undergoing collective cell migration after treatment by LY292004 compound. 1 V/cm EF (the cathode at the right) was applied for 30 min. Note the dark blue regions at the groups’ peripheries, indicating transient movements of cells at the groups’ peripheries to the anode. D.Comparison of the time series of the spatially averaged directedness of the group’s front one/third (green), center one/third (black) and rear one/third (blue) demonstrates that all cells within the group respond to EF directionally on the same time scale. Three parts of the group were defined by two vertical lines separating the disc-like cell group into three parts with equal areas. E.A peripheral group of cells splits from the large cell group undergoing EF-directed collective cell migration after treatment with 100 µm of LY292004 compound. Times are in hh:mm:ss before and after applying 1 V/cm EF (the cathode is at the right). Red arrow shows the trajectory of the centroid of the keratocyte group, while yellow arrow shows the trajectory of the centroid of the splitting keratocyte group. Bar, 200 µm. F. Computational result: overlays of a representative PI3K-inhibited cell group with a long narrow appendix in the beginning (gray) and end (magenta) of the simulation. The snapshots are taken at times 50 and time 330 (time is not physical; measured in computational steps). The result is that the appendix sub-group splits from the large body of the cell group, consistent with the experimental result shown in (E).

**Supplemental Table:**
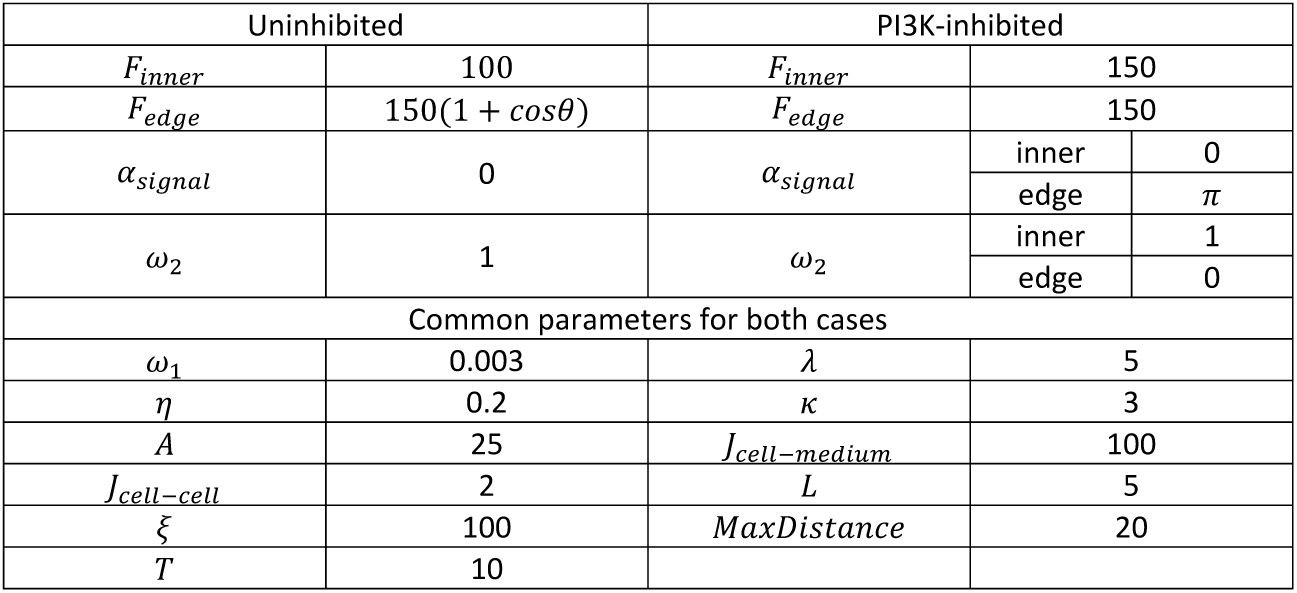
model parameters.

## REFERENCES

1. Friedl, P. and D. Gilmour, Collective cell migration in morphogenesis, regeneration and cancer. Nat Rev Mol Cell Biol, 2009. 10(7): p. 445–57.

2. Scarpa, E. and R. Mayor, Collective cell migration in development. J Cell Biol, 2016. 212(2): p. 143–55.

3. Szabo, B., et al., Phase transition in the collective migration of tissue cells: experiment and model. Phys Rev E Stat Nonlin Soft Matter Phys, 2006. 74(6 Pt 1): p. 061908.

4. Schumacher, L.J., et al., Multidisciplinary approaches to understanding collective cell migration in developmental biology. Open Biol, 2016. 6(6).

5. Camley, B.A., et al., Emergent Collective Chemotaxis without Single-Cell Gradient Sensing. Phys Rev Lett, 2016. 116(9): p. 098101.

6. Copenhagen, K., et al., Frustration-induced phases in migrating cell clusters. Sci Adv, 2018. 4(9): p. eaar8483.

7. Omelchenko, T., et al., Rho-dependent formation of epithelial “leader” cells during wound healing. Proc Natl Acad Sci U S A, 2003. 100(19): p. 10788–93.

8. Reffay, M., et al., Interplay of RhoA and mechanical forces in collective cell migration driven by leader cells. Nat Cell Biol, 2014. 16(3): p. 217–23.

9. Cohen, D.J., W.J. Nelson, and M.M. Maharbiz, Galvanotactic control of collective cell migration in epithelial monolayers. Nat Mater, 2014. 13(4): p. 409–17.

10. Trepat, X. and J.J. Fredberg, Plithotaxis and emergent dynamics in collective cellular migration. Trends Cell Biol, 2011. 21(11): p. 638–46.

11. Shellard, A., et al., Supracellular contraction at the rear of neural crest cell groups drives collective chemotaxis. Science, 2018. 362(6412): p. 339–343.

12. Levin, M., G. Pezzulo, and J.M. Finkelstein, Endogenous Bioelectric Signaling Networks: Exploiting Voltage Gradients for Control of Growth and Form. Annu Rev Biomed Eng, 2017. 19: p. 353–387.

13. Sun, Y., et al., Infection-generated electric field in gut epithelium drives bidirectional migration of macrophages. PLoS Biol, 2019. 17(4): p. e3000044.

14. Ferreira, F., et al., Early bioelectric activities mediate redox-modulated regeneration. Development, 2016. 143(24): p. 4582–4594.

15. Guo, A., et al., Effects of physiological electric fields on migration of human dermal fibroblasts. J Invest Dermatol, 2010. 130(9): p. 2320–7.

16. Chang, F. and N. Minc, Electrochemical control of cell and tissue polarity. Annu Rev Cell Dev Biol, 2014. 30: p. 317–36.

17. Li, L., et al., E-cadherin plays an essential role in collective directional migration of large epithelial sheets. Cell Mol Life Sci, 2012. 69(16): p. 2779–89.

18. Lalli, M.L. and A.R. Asthagiri, Collective migration exhibits greater sensitivity but slower dynamics of alignment to applied electric fields. Cell Mol Bioeng, 2015. 8(2): p. 247–257.

19. Zajdel, T.J., et al., SCHEEPDOG: Programming Electric Cues to Dynamically Herd Large-Scale Cell Migration. Cell Syst, 2020. 10(6): p. 506-514.e3.

20. Zhang, Y., et al., Collective cell migration has distinct directionality and speed dynamics. Cell Mol Life Sci, 2017. 74(20): p. 3841–3850.

21. Cho, Y., et al., Electric field-induced migration and intercellular stress alignment in a collective epithelial monolayer. Mol Biol Cell, 2018. 29(19): p. 2292–2302.

22. Mogilner, A., E.L. Barnhart, and K. Keren, Experiment, theory, and the keratocyte: An ode to a simple model for cell motility. Semin Cell Dev Biol, 2019.

23. Cooper, M.S. and M. Schliwa, Motility of cultured fish epidermal cells in the presence and absence of direct current electric fields. J Cell Biol, 1986. 102(4): p. 1384–99.

24. Allen, G.M., A. Mogilner, and J.A. Theriot, Electrophoresis of cellular membrane components creates the directional cue guiding keratocyte galvanotaxis. Curr Biol, 2013. 23(7): p. 560–8.

25. Sun, Y., et al., Keratocyte fragments and cells utilize competing pathways to move in opposite directions in an electric field. Curr Biol, 2013. 23(7): p. 569–74.

26. Rapanan, J.L., et al., Collective cell migration of primary zebrafish keratocytes. Exp Cell Res, 2014. 326(1): p. 155–65.

27. Huang, L., et al., The involvement of Ca2+ and integrins in directional responses of zebrafish keratocytes to electric fields. J Cell Physiol, 2009. 219(1): p. 162–72.

28. Sun, Y.H., et al., An Experimental Model for Simultaneous Study of Migration of Cell Fragments, Single Cells, and Cell Sheets. Methods Mol Biol, 2016. 1407: p. 251–72.

29. Sun, Y.H., et al., Electric fields accelerate cell polarization and bypass myosin action in motility initiation. J Cell Physiol, 2018. 233(3): p. 2378–85.

30. Song, B., et al., Application of direct current electric fields to cells and tissues in vitro and modulation of wound electric field in vivo. Nat Protoc, 2007. 2(6): p. 1479–89.

31. Zhao, M., et al., Directed migration of corneal epithelial sheets in physiological electric field. Invest Ophthalmol Vis Sci, 1996. 37(13): p. 2548–58.

32. Nakajima, K., et al., KCNJ15/Kir4.2 couples with polyamines to sense weak extracellular electric fields in galvanotaxis. Nat Commun, 2015. 6.

33. Sun, Y.H., et al., Airway epithelial wounds in rhesus monkey generate ionic currents that guide cell migration to promote healing. J Appl Physiol (1985), 2011. 111(4): p. 1031–41.

34. Gruler, H. and R. Nuccitelli, Neural crest cell galvanotaxis: new data and a novel approach to the analysis of both galvanotaxis and chemotaxis. Cell Motil Cytoskeleton, 1991. 19(2): p. 121–33.

35. Tai, G., et al., Electrotaxis and wound healing: experimental methods to study electric fields as a directional signal for cell migration. Methods Mol Biol, 2009. 571: p. 77–97.

36. Pincus, Z. and J.A. Theriot, Comparison of quantitative methods for cell-shape analysis. J Microsc, 2007. 227(Pt 2): p. 140–56.

37. Barnhart, E.L., et al., An adhesion-dependent switch between mechanisms that determine motile cell shape. PLoS Biol, 2011. 9(5): p. e1001059.

38. Keren, K., et al., Mechanism of shape determination in motile cells. Nature, 2008. 453(7194): p. 475–80.

39. Graner, F. and J.A. Glazier, Simulation of biological cell sorting using a two-dimensional extended Potts model. Phys Rev Lett, 1992. 69(13): p. 2013–2016.

40. Swat, M.H., et al., Multi-scale modeling of tissues using CompuCell3D. Methods Cell Biol, 2012. 110: p. 325–66.

41. McCaig, C.D., B. Song, and A.M. Rajnicek, Electrical dimensions in cell science. J Cell Sci, 2009. 122(Pt 23): p. 4267–76.

42. Angelini, T.E., et al., Cell migration driven by cooperative substrate deformation patterns. Phys Rev Lett, 2010. 104(16): p. 168104.

43. Vedula, S.R., et al., Emerging modes of collective cell migration induced by geometrical constraints. Proc Natl Acad Sci U S A, 2012. 109(32): p. 12974–9.

44. Szabo, A., et al., In vivo confinement promotes collective migration of neural crest cells. J Cell Biol, 2016. 213(5): p. 543–55.

45. Varennes, J., B. Han, and A. Mugler, Collective Chemotaxis through Noisy Multicellular Gradient Sensing. Biophys J, 2016. 111(3): p. 640–649.

46. Malet-Engra, G., et al., Collective cell motility promotes chemotactic prowess and resistance to chemorepulsion. Curr Biol, 2015. 25(2): p. 242–250.

47. Camley, B.A., Collective gradient sensing and chemotaxis: modeling and recent developments. J Phys Condens Matter, 2018. 30(22): p. 223001.

48. Efimova, N. and T.M. Svitkina, Branched actin networks push against each other at adherens junctions to maintain cell-cell adhesion. J Cell Biol, 2018. 217(5): p. 1827–1845.

49. Anon, E., et al., Cell crawling mediates collective cell migration to close undamaged epithelial gaps. Proc Natl Acad Sci U S A, 2012. 109(27): p. 10891–6.

50. Lee, P. and C.W. Wolgemuth, Crawling cells can close wounds without purse strings or signaling. PLoS Comput Biol, 2011. 7(3): p. e1002007.

51. Mertz, A.F., et al., Cadherin-based intercellular adhesions organize epithelial cell-matrix traction forces. Proc Natl Acad Sci U S A, 2013. 110(3): p. 842–7.

52. Fortuna, I., et al., CompuCell3D Simulations Reproduce Mesenchymal Cell Migration on Flat Substrates. Biophys J, 2020. 118(11): p. 2801–2815.

53. Rens, E.G. and L. Edelstein-Keshet, From energy to cellular forces in the Cellular Potts Model: An algorithmic approach. PLoS Comput Biol, 2019. 15(12): p. e1007459.

54. Xu, J., et al., Divergent signals and cytoskeletal assemblies regulate self-organizing polarity in neutrophils. Cell, 2003. 114(2): p. 201–14.

55. Yamaguchi, N., et al., Leader cells regulate collective cell migration via Rac activation in the downstream signaling of integrin beta1 and PI3K. Sci Rep, 2015. 5: p. 7656.

56. Durant, F., et al., The Role of Early Bioelectric Signals in the Regeneration of Planarian Anterior/Posterior Polarity. Biophys J, 2019. 116(5): p. 948–961.

57. Farooqui, R. and G. Fenteany, Multiple rows of cells behind an epithelial wound edge extend cryptic lamellipodia to collectively drive cell-sheet movement. J Cell Sci, 2005. 118(Pt 1): p. 51–63.

58. Etienne-Manneville, S., Neighborly relations during collective migration. Curr Opin Cell Biol, 2014. 30: p. 51–9.

59. Sato, M.J., et al., Switching direction in electric-signal-induced cell migration by cyclic guanosine monophosphate and phosphatidylinositol signaling. Proc Natl Acad Sci U S A, 2009. 106(16): p. 6667–72.

60. Camley, B.A. and W.J. Rappel, Cell-to-cell variation sets a tissue-rheology-dependent bound on collective gradient sensing. Proc Natl Acad Sci U S A, 2017. 114(47): p. E10074–e10082.

